# Thr50 O-GlcNAcylated PKM2 promotes aerobic glycolysis and pancreatic ductal adenocarcinoma progression via ARNT-mediated CDC27 transnational activation of the AKT pathway

**DOI:** 10.64898/2026.07.27.740965

**Authors:** Bin Yang, Yi Zhu, Xiaopeng Zhan, Yun Zhang, Jianing Cui, Yu Zhou, Shikai Zhu

## Abstract

Pancreatic ductal adenocarcinoma (PDAC) is one of the most lethal cancers and more evidence suggests that glucose metabolism plays a significant role in the development and progression with glycosylation at multiple sites potentially being a key characteristic. However, the underlying mechanisms remain insufficiently studied. Here, we first confirmed the presence of O-GlcNAc glycosylation modifications at Thr50 on PKM, which is highly expressed in PDAC and strongly correlated with poor prognosis. Further, we found that PKM expression was significantly positively correlated with CDC27 levels, and mutation of the Thr50 O-GlcNAc site in PKM abolished the upregulation of CDC27. We confirmed that O-GlcNAc-modified PKM enhances nuclear translocation of ARNT, which binds to the CDC27 promoter to upregulate its expression. Finally, we demonstrated that reduced expression of CDC27, as a key component of the APC/C complex, leads to downregulation of ubiquitination at the K11 site of PPP2CA, resulting in upregulation of PPP2CA expression, in turn, reduces AKT phosphorylation, ultimately driving PDAC regression by inhibiting aerobic glycolysis. Thus, we delineate a novel O-GlcNAcylation-dependent pathway where PKM drives PDAC progression through ARNT-mediated CDC27 transcriptional activation and AKT-mediated glycolysis.

## Introduction

Pancreatic ductal adenocarcinoma (PDAC) is an aggressive malignancy with limited therapeutic options(1), exhibiting rising global incidence with mortality-to-incidence ratio approaching. Epidemiological data (2015–2023) show a >50% increase in PDAC-related mortality(2). Current therapeutic strategies remain palliative, with only 15% of patients qualifying for curative resection at diagnosis and systemic therapies showing <40% objective response rates(3). Emerging evidence implicates metabolic reprogramming—particularly enhanced aerobic glycolysis—as a hallmark of PDAC pathogenesis(4).

The pyruvate kinase M (PKM) isoforms (PKM1/PKM2), key regulators of glycolytic flux, undergo dynamic tetramer-dimer transitions that determine their catalytic activity and metabolic plasticity(5). O-GlcNAc transferase (OGT)-mediated post-translational modification of PKM2 at Thr405/Ser406 stabilizes its dimeric form, diverting glycolytic intermediates toward anabolic pathways to fuel tumor proliferation(6).

O-GlcNAcylation—a nutrient-sensitive modification involving β-linked N-acetylglucosamine attachment to Ser/Thr residues—is aberrantly elevated in PDAC (7). Cancer cells shunt ∼3-5% of glycolytic flux into the hexosamine biosynthesis pathway (HBP) to sustain O-GlcNAcylation dynamics(8). This metabolic rewiring enables PDAC cells to bypass oxidative phosphorylation while generating biosynthetic precursors through aerobic glycolysis(9).

Cell Division Cycle 27 (CDC27), an essential component of the anaphase-promoting complex/cyclosome (APC/C), exhibits context-dependent roles in tumorigenesis (10). Current research suggests that CDC27 may play significant roles in apoptosis, stemness, endocytosis, and epithelial-mesenchymal transition (EMT). However, due to the complexity and diversity of its mechanisms, its functions vary across different cancers. For instance, While CDC27 promotes EMT and metastasis in colorectal cancer(11), its downregulation confers chemoresistance in gliomas through distinct mechanisms(12). Consequently, CDC27 cannot be straightforwardly categorized as either an oncogene (OG) or a tumor suppressor gene (TSG). Given the distinctive role of CDC27, it is imperative to investigate its function in PDAC.

This study reveals that O-GlcNAcylation of PKM at Thr50 enhances glycolytic flux through ARNT-mediated transcriptional upregulation of CDC27, which subsequently activates AKT signaling. This axis ultimately drives the progression of PDAC and thus may be an effective targeted therapeutic approach.

## Material and methods

### Clinical Sample Collection

Formalin-fixed paraffin-embedded (FFPE) PDAC tumor tissues and adjacent normal pancreatic tissues were obtained from a tissue microarray (HPanAde180Sur-02, Shanghai Outdo Biotech Co., Ltd., China). All samples were collected with written informed consent from patients, and the study was approved by the Institutional Review Board of the Second Affiliated Hospital, Zhejiang University (Ethics Committee Approval No. [AIRB-2022-1476]). The use of human tissues complied with the Declaration of Helsinki and relevant ethical guidelines.

### Immunohistochemistry (IHC) staining

The tissues were dewaxed and subjected to peroxidase blocking. Then, the tissues were subjected to antigen retrieval, primary antibody incubation (anti-PKM2, 1:150, #PB9379, Biological Technology; anti-CDC27, 1:150, #10918-1-AP, Proteintech Group) overnight at 4°C in a humidified chamber. After washing (3 × 5 min with PBS), biotinylated goat anti-rabbit secondary antibody (1:200, Vector Laboratories, USA) was added for 1 h at RT. Signal amplification was achieved using the ABC Elite Kit (Vector Laboratories), followed by visualization with diaminobenzidine (DAB) chromogen (Dako, Denmark) for 5 min. Slides were counterstained with hematoxylin (Sigma-Aldrich, USA), dehydrated through ethanol, cleared in xylene, and mounted with Entellan® mounting medium (Merck KGaA, Germany).

### Cell culture

Human PANC-1, PATU 8849s cells were purchased from the Cell Bank of the Chinese Academy of Science. Cells were cultured in regular 1640 or DMEM medium plus 10% FBS and 1% penicillin/streptomycin. All cells were grown at 37°C in the presence of 5% CO_2_. Testing for mycoplasma contamination is carried out every six months and STR profiles of the above cells are tested.

### Stable cell lines construction

Three shRNA sequences targeting human PKM2 (shPKM2-1: AGGGAAAGAACATCAAGATTA; shPKM2-2: TGGATAACGCCTACATGGAAA; shPKM2-3: CAGCAAGATCTACGTGGATGA) and shRNA sequences targeting CDC27 (shCDC27-1: GAGGTTAGAAGGATTGAGAAT; shCDC27-2: CACTACAATACTGGTTGGGTA; shCDC27-3: TCCCAAAGAATCCCTCGTTTA) were designed and cloned into the lentiviral vector BR-V108 (GenScript, Nanjing, China) between EcoR I (#R0101L, New England Biolabs, USA) and Age I (#R3552L, New England Biolabs, USA) restriction sites. The scrambled sequence (TTCTCCGAACGTGTCACGT) was used as a negative control. For PKM2 overexpression, a cDNA fragment encoding full-length human PKM2 was amplified by PCR and ligated into the same vector.

Lentiviral particles were produced in HEK293T cells using Lipofectamine 3000 (Thermo Fisher Scientific, USA) according to the manufacturer’s protocol. Briefly, HEK293T cells were co-transfected with the shRNA/overexpression vectors, packaging plasmid psPAX2 (Addgene, USA), and envelope plasmid pMD2.G (Addgene, USA). Viral supernatants collected 72 hours post-transfection, concentrated using PEG-it Virus Concentration Solution (System Biosciences, USA), and stored at −80°C. For transduction, PANC-1 and PATU 8849s cells were infected with lentivirus in the presence of 8 μg/mL polybrene (Sigma-Aldrich, USA). Stable cell lines were selected with puromycin (0.5 μg/mL for PANC-1, 0.8 μg/mL for PATU 8849s; MedChemExpress, USA) for 7 days.

### Quantitative real-time PCR (qPCR)

Total RNA in cells was extracted using TRIzol RNA isolation system (Invitrogen) and converted to cDNA using PrimeScript RT Reagent Kit (TaKaRa). Then q-PCR was performed with a 7500 Fast™ System (Applied Biosystems) using the Sensi Mix SYBR Kit (Bio-Rad). The mRNA level was calculated using the 2-ΔΔCt method and normalized to GAPDH. The primers sequences are as below. PKM2: CATGGCTCCTACGGAGAGGT and ACATGGAACGCTTTACCGCAT; CDC27: AAACCGCACCAAAAGCCAC and ATCCTTGGGTTCACTCGTCTT; GAPDH: TGACTTCAACAGCGACACCCA and CATGGGCCACGATCCTCTTTA.

### Chromatin immunoprecipitation (ChIP)

Cells were treated with plasmids for 48 h, then CHIP was performed using Pierce Agarose ChIP Kit (No.26156, Thermo Fisher Scientific). DNA template enrichment was analysed by qPCR using primers specific to each target genes promoter. The primers sequences are as below. CDC27-ARNT-F1: GCTTCATAATCTACAGCACACCAA and TTCTCAGCCACGCCTTCAG; CDC27-ARNT-F2: GCTTCATAATCTACAGCACACCAA and CTACGCCAAATCTGCAACCA; CDC27-ARNT-F3: TTGTCAACCACCAAGTCAAGAA and AGATTCAAGATGGCTTCACTCAC.

### Western blot analysis

Cells were lysed in RIPA lysis buffer (Sigma) containing phosphorylase and protease inhibitors (Sigma) and the proteins were separated through electrophoresis in 8–15% SDS-PAGE gels. Then the proteins were transferred from the gels to PVDF (polyvinylidene difluoride) membranes (Bio-Rad) and incubated with the indicated antibodies (PKM, 1:1000, #10078-2-AP, Proteintech; HK2, 1:3000, #66974-1-Ig, Proteintech; CDC27, 1:4000, #10918-1-AP, Proteintech; PGLUT1, 1:1000, #21829-1-AP, Proteintech; LDHA, 1:3000, #ab52488, Abcam; ARNT, 1:1000, #sc-17811, Santa Cruz; Histone H3, 17168-1-AP, #Proteintech; K11, #51064-2-AP, Proteintech; PPP2CA, #13482-1-AP, Proteintech; AKT, 1:3000, #10176-2-AP, Proteintech; p-AKT, 1:5000, #66444-1-Ig, Proteintech; GAPDH, 1:30000, #60004-1-lg, Proteintech; β-Actin, 1:4000, #66009-1-Ig, Proteintech). After incubation with a secondary antibody (Goat Anti-Rabbit, 1:3000, #A0208, Beyotime; Goat Anti-Mouse, 1:3000, #A0216, Beyotime), the proteins were visualized by chemiluminescence. All original western blots are shown in Supplementary original data.

### Immunoprecipitation (IP) and Co-immunoprecipitation (CO-IP)

Cells were lysed in Pierce IP buffer (Thermo Fisher) containing protease and phosphatase inhibitors (Sigma). After incubation with indicated antibodies, the lysates were mixed with protein A/G agarose (Thermo Fisher). For proteins with tag, we used immunomagnetic beads coupled with anti-Flag/anti-HA antibody. After IP, protein A/G agarose or magnetic beads were washed three times with TBST (0.1% Tween-20, 150 mM NaCl, 10 mM Tris-HCl pH7.5) and eluted in SDS lysis buffer (100 mM NaCl, 1%SDS, 50 mM Tris-HCl pH 7.5) for western blots.

### Cell proliferation assays

For CCK-8 proliferation assay, cells were seeded into 96-well plates at a density of 3,000 cells/well in 100 μL of complete medium supplemented with 10% FBS. After 24 h of incubation at 37°C with 5% CO_2_, cells were treated with varying concentrations of experimental agents. At specified time points (24, 48, 72, 96 and 120 h), 10 μL of CCK-8 reagent was added to each well and incubated for 1–4 h. Absorbance at 450 nm was measured using a microplate reader (SpectraMax iD5, Molecular Devices, USA).

For plate clone formation assay, cells were trypsinized, resuspended in complete medium, and serially diluted to final densities of 50, 100, and 200 cells/well in 6-well plates (Corning, USA, #353046). Plates were incubated at 37°C for 14–21 days until visible colonies formed (>50 cells/colony). Cultures were fixed with 4% paraformaldehyde (Sigma-Aldrich, USA, #P6148) for 15 min and stained with 0.1% crystal violet (Sigma-Aldrich, #C0775) for 30 min. Colonies were imaged and counted manually.

### Annexin V-PI double staining and flow cytometry analysis

Apoptosis was quantified using the Annexin V-FITC/Propidium Iodide (PI) double-staining assay (BD Pharmingen, USA, #556547) following optimized protocols. Cells were gently dissociated using 0.25% Trypsin-EDTA (Gibco, USA, #25300-054) for ≤3 min at 37°C to minimize membrane damage. Cell suspensions were centrifuged at 300 × g for 5 min, washed twice with ice-cold PBS (Thermo Fisher Scientific, USA, #10010023), and resuspended in 1× Annexin V Binding Buffer (BD Pharmingen, #556454) at 1 × 10^6^ cells/mL. 100 μL of cell suspension was transferred to flow cytometry tubes (BD Falcon, USA, #352058). 5 μL of Annexin V-FITC and 5 μL of PI (final concentration: 1 μg/mL each) were added, mixed gently, and incubated in the dark at 25°C for 15 min. 400 μL of Annexin V Binding Buffer was added to stop the reaction, and samples were analyzed within 1 h to prevent fluorescence quenching. Data was acquired on a BD FACSCanto II (BD Biosciences, USA) equipped with 488 nm (FITC/PI) and 633 nm (PE-Cy7) lasers. Apoptosis rates were calculated as: Apoptosis Rate (%) = (Q1-UR + Q1-LR)/Total Cells×100%.

### Cell migration assays

For wound healing assay, cells were seeded into 96-well plates at a density of 5 × 10^5^ cells/well in DMEM medium supplemented. After 24 h of incubation, confluent monolayers were scratched using a 200 μL sterile pipette tip (BD Falcon, #352058) perpendicular to pre-marked reference lines on the plate’s underside. Three parallel scratches were made per well to ensure consistency. Non-migrated cells and debris were removed by gently washing three times with PBS (Thermo Fisher Scientific, #10010023). Cells were cultured in serum-free medium (RPMI-1640 + 0.1% BSA) to minimize proliferation interference. At 0, 24, and 48 h post-scratch, phase-contrast images were captured using an inverted microscope (Olympus IX73) at ×10 magnification. The percentage of wound closure was calculated as: Wound Closure = (Initial Wound Width−Wound Width at Time t)/Initial Wound Width.

For Transwell assay, cell migratory capacity was further assessed using Transwell chambers (Corning, #3422) with 8 μm pore membranes. Cells (1 × 10^4^ per well) were resuspended in serum-free medium and loaded into the upper chamber. The lower chamber contained 600 μL of complete medium with 10% FBS as a chemoattractant. Non-migrated cells on the upper membrane were removed using a cotton swab (BD Falcon, #352058). Migrated cells were fixed with 4% paraformaldehyde (Sigma-Aldrich, #P6148) for 15 min and stained with 0.1% crystal violet (Sigma-Aldrich, #C0775) for 30 min. Membranes were imaged (×20 magnification), and cells were counted manually. Migration rate was expressed as cells per high-power field (HPF).

### Glycolytic parameter measurements

For glucose content assay, intracellular glucose levels were quantified using the Glucose Uptake-Glo^TM^ Assay (Promega, #J1341) following the manufacturer’s protocol. Briefly, cells were seeded in 96-well plates (Corning, #3599) at 5 × 10^4^ cells/well and cultured for 24 h. After treatment, 50 μL of glucose detection reagent was added, followed by a 30-min incubation at room temperature. Luminescence was measured using a microplate reader (SpectraMax iD5, Molecular Devices). Glucose uptake rates were calculated based on standard curves generated with glucose standards (Promega, #G7521).

ATP levels were measured using the EnzyLight ATP Assay Kit (BioAssay Systems, #EATP-048). Cells (1 × 10^4^/well) were lysed in 50 μL of ATP assay buffer, and 50 μL of luciferase reagent was added. Luminescence was immediately recorded using a microplate reader. ATP concentrations were determined using a standard curve (0.1–10 μM) provided with the kit.

Extracellular lactate was quantified using the L-Lactate Assay Kit (Megazyme, #K-LATE). Cell culture supernatants (50 μL) were mixed with 50 μL of assay buffer and 10 μL of lactate dehydrogenase reagent. The reaction was incubated at 37°C for 15 min, and absorbance at 340 nm was measured. Lactate concentrations were calculated using a standard curve (0.2–20 mM).

Extracellular Acidification Rate (ECAR) and Oxygen Consumption Rate (OCR) were measured using the Seahorse XF Real-Time ATP Rate Assay Kit (Agilent, #103675-100) on a Seahorse XF96 Analyzer (Agilent). Cells (2 × 10^4^/well) were plated in XF96 cell culture plates (Agilent, #101085-004) and allowed to equilibrate for 1 h. Basal ECAR and OCR were recorded, followed by sequential injections of 1 μM oligomycin, 1 μM rotenone/antimycin A, and 50 mM 2-deoxyglucose (2-DG). Data was analyzed using Seahorse Wave software (Agilent) to calculate glycolytic and mitochondrial ATP production rates.

### Tumour growth experiments in vivo

All animal experiments were approved by the ethics committee of Zhejiang University. For the xenograft assay, male athymic nude mice (6-8 weeks), obtained from the SLAC (Shanghai SLAC Laboratory Animal Co., Ltd.), were randomly divided into 3 groups. A total of 1 × 106 cells in 100 µL PBS were injected into the right subcutaneous of nude mice. The growth of tumours was observed every 3 days. At 20 days after the injection, all mice were sacrificed, and the tumours were harvested to detect tumour volume (Vtumour = 0.5 × L × W^2^; L = length, W = width). Then the tumours were paraffin-embedded and sectioned for HE (hematoxylin-eosin) and Ki67 staining. The investigators were blinded to the group allocation during the experiment and when assessing the outcome.

## Statistical analysis

All experiments were performed in triplicates. All data are shown as the mean ± SD. Data were analyzed using two-tailed t-tests, Chi-square (x2) tests or one-way analysis of variance (ANOVA). Kaplan–Meier survival analyses were used to compare survival times among PDAC patients based on gene expression; the log-rank test was used to generate P values. Cox proportional hazards regression analyses were used to assess the effect of clinical variables on patient survival. Univariate and multivariate analyses were used to assess the influence of clinical variables on survival. The p values and hazard ratios are indicated. P values < 0.05 were considered statistically significant. (*P < 0.05; **P < 0.01; ***P < 0.001; ****P < 0.0001). All histograms and curves were constructed with GraphPad Prism 8.0.2 software (GraphPad Software, La Jolla, CA, USA) and SPSS V.21.0 software (SPSS, Inc., Chicago, IL, USA).

## Results

### O-GlcNAcylated PKM Predicts PDAC Adverse Clinical Outcomes

To delineate the role of O-GlcNAcylation in pancreatic carcinogenesis, we conducted immunoprecipitation-mass spectrometry (IP-MS) in PATU8988S cells, identifying seven O-GlcNAcylated proteins including PKM, CFL1, and HSP90AA1 (**Figure 1A**). Subsequently, lentiviral shRNA-mediated silencing in PANC-1 cells revealed that PKM depletion caused the most profound proliferation inhibition (p<0.001), exceeding the effects of EEF2 and RPS2 knockdown (**Figure 1B**). These data establish PKM as a key O-GlcNAc-modified oncoprotein in PDAC.

**Figure 1.**
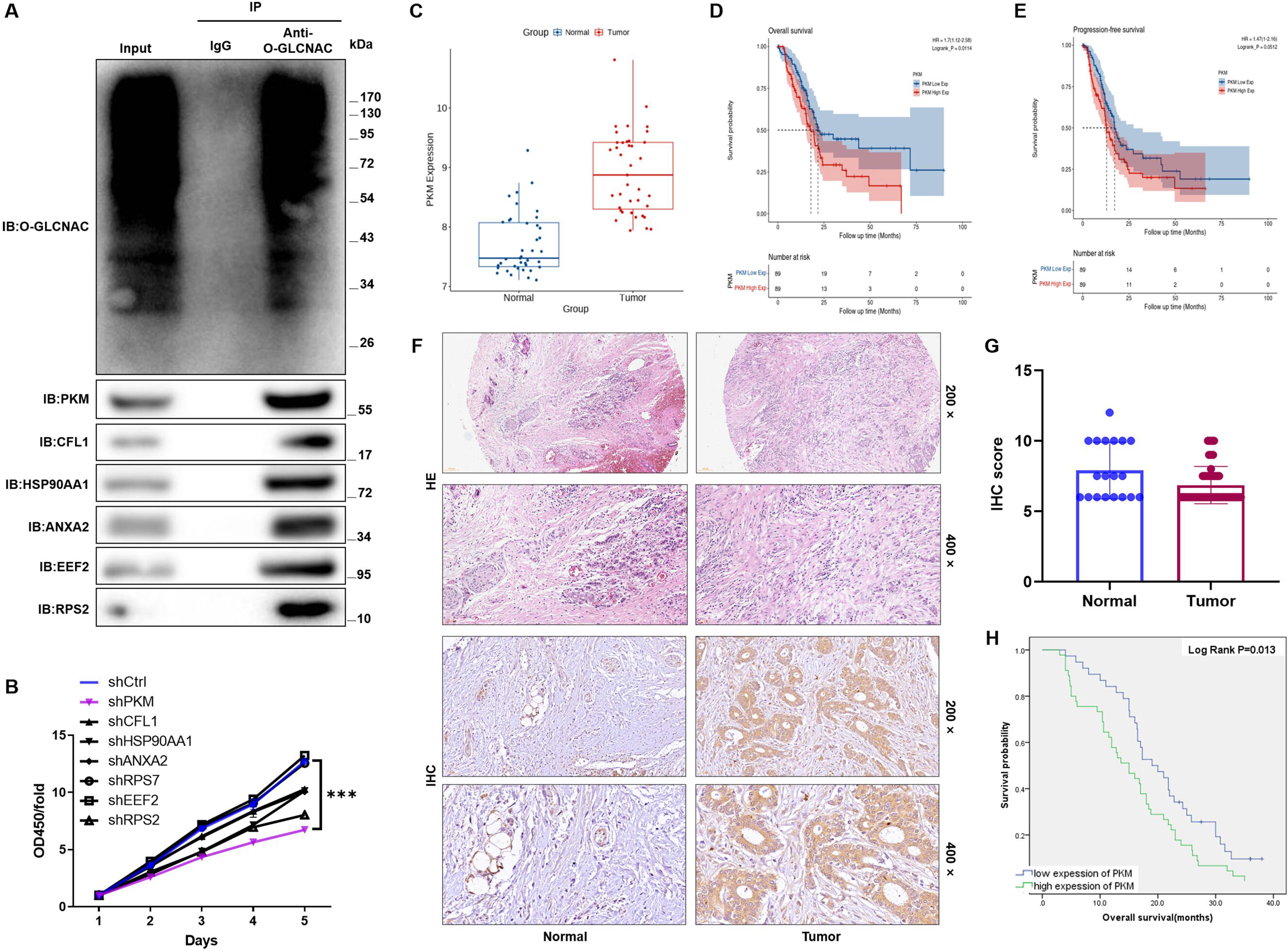
O-GlcNAcylation of PKM predicts adverse clinical outcomes in PDAC. **(A)** Immunoprecipitation-mass spectrometry (IP-MS) identified seven O-GlcNAcylated proteins in PATU8988S cells, including PKM, CFL1, and HSP90AA1. **(B)** Lentiviral shRNA-mediated knockdown in PANC-1 cells revealed that PKM depletion significantly inhibited cell proliferation compared to EEF2 or RPS2 knockdown (p < 0.001). **(C)** Public datasets GSE15471 confirmed PKM mRNA upregulation in PDAC tumors versus normal tissues (p < 0.001). (D-E) Kaplan-Meier analyses demonstrated that high PKM expression correlated with reduced overall survival **(D)** (HR=1.7, 95% CI 1.12-2.58; log-rank p=0.0114) and progression-free survival **(E)** (HR=1.47, 95% CI 1-2.16; p=0.0512). **(F-G)** Representative hematoxylin-eosin (HE) staining and immunohistochemical (IHC) analysis **(F)** of PKM expression in clinical PDAC and normal pancreatic tissues. **(G)** H-score quantification revealed significantly elevated PKM levels in tumors (p < 0.001). **(H)** Kaplan-Meier survival curves stratified by PKM IHC scores in a clinical cohort showed shorter overall survival in patients with high PKM expression (p < 0.001).

Interrogation of the GSE15471 dataset (tumors vs. normals) confirmed PKM mRNA upregulation in PDAC (p<0.001; (**Figure 1C**). Kaplan-Meier analysis demonstrated that high PKM expression correlated with reduced median overall survival (HR=1.7, 95% CI 1.12-2.58; log-rank p=0.0114) (**Figure 1D**) and progression-free survival (HR=1.47, 95% CI 1-2.16; p=0.0512) (**Figure 1E**).

To further validate these findings, we constructed tissue microarrays (TMAs) using clinical samples from normal and tumor tissues, followed by hematoxylin-eosin (HE) staining and immunohistochemical (IHC) assays. The histopathological staining (**Figure 1F**)combined with H-score analysis **(Figure 1G**; **Table 1)** confirmed significantly elevated PKM expression in PDAC tissues. Moreover, Kaplan-Meier analysis based on clinical follow-up data demonstrated that higher PKM expression correlated with shorter patient survival (**Figure 1H**).

**Table 1.** H-score analysis of tissue microarrays using clinical samples from normal and tumor tissues, with HE staining and IHC assays.

| PKM<br>expression | Tumor tissue |  | Para-carcinoma tissue |  | p value |
| --- | --- | --- | --- | --- | --- |
|  | Cases | Percentage | Cases | Percentage |  |
| Low | 38 | 45.8% | 61 | 76.3% | 0.005 |
| High | 45 | 54.2% | 19 | 23.7% |  |

To elucidate the correlation between PKM expression levels and clinicopathological characteristics in PDAC, we performed Mann-Whitney U tests, which revealed that PKM expression levels showed significant positive correlations with age (p=0.037), stage (p=0.030), metastasis (p=0.020), tumor differentiation grade (p=0.009) **(Table 2)**. Spearman’s rank correlation coefficients confirmed positive associations with age (ρ=0.231, p=0.036), stage (ρ=0.239, p=0.029), and metastasis (ρ=0.257, p=0.019), but inverse correlation with differentiation grade (ρ=-0.290, p=0.008) **(Table 3)**. These data establish that O-GlcNAc-modified PKM serves as both a metabolic driver and prognostic biomarker in PDAC, with its overexpression mechanistically linked to tumor progression and dismal clinical outcomes.

**Table 2.** Mann-Whitney U analysis of PKM expression correlation with PDAC clinicopathological characteristics.

| Features | No. of patients | PKM expression |  | p value |
| --- | --- | --- | --- | --- |
|  |  | low | high |  |
| All patients | 83 | 38 | 45 |  |
| Age (years) |  |  |  | 0.037 |
| < 60 | 42 | 24 | 18 |  |
| ≥60 | 41 | 14 | 27 |  |
| Gender |  |  |  | 0.345 |
| Male | 50 | 25 | 25 |  |
| Female | 33 | 13 | 20 |  |
| Stage |  |  |  | 0.030 |
| I | 24 | 12 | 12 |  |
| II | 27 | 17 | 10 |  |
| III | 10 | 5 | 5 |  |
| IV | 22 | 4 | 18 |  |
| T Infiltrate |  |  |  | 0.713 |
| T1 | 6 | 3 | 3 |  |
| T2 | 35 | 17 | 18 |  |
| T3 | 32 | 13 | 19 |  |
| T4 | 10 | 5 | 5 |  |
| Metastasis |  |  |  | 0.020 |
| M0 | 62 | 33 | 29 |  |
| M1 | 21 | 5 | 16 |  |
| lymphatic metastasis (N) |  |  |  | 0.948 |
| N0 | 30 | 15 | 15 |  |
| N1 | 42 | 16 | 26 |  |
| N2 | 11 | 7 | 4 |  |
| Degree of differentiation |  |  |  | 0.009 |
| Low | 4 | 0 | 4 |  |
| Medium | 70 | 31 | 39 |  |
| High | 9 | 7 | 2 |  |
| Surgical margin R |  |  |  | 0.217 |
| R0 | 75 | 36 | 39 |  |
| R1 | 8 | 2 | 6 |  |
| Vascular invasion |  |  |  | 0.819 |
| NO | 47 | 21 | 26 |  |
| YES | 36 | 17 | 19 |  |
| perineural invasion |  |  |  | 0.270 |
| NO | 27 | 10 | 17 |  |
| YES | 56 | 28 | 28 |  |

**Table 3.** Spearman correlation analysis of PKM expression and PDAC clinicopathological indicators.

|  |  | PKM |
| --- | --- | --- |
| Age | Spearman's correlationan | 0.231 |
|  | Two-tailed Test | 0.036 |
|  | N | 83 |
| Stage | Spearman's correlationan | 0.239 |
|  | Two-tailed Test | 0.029 |
|  | N | 83 |
| Metastasis | Spearman's correlationan | 0.257 |
|  | Two-tailed Test | 0.019 |
|  | N | 83 |
| Degree of differentiation | Spearman's correlationan | -0.290 |
|  | Two-tailed Test | 0.008 |
|  | N | 83 |

### Knockdown of O-GlcNAcylated PKM Inhibits the Proliferation and Migration of PDAC Cells

To delineate the functional role of PKM in pancreatic oncogenesis, we generated stable PKM-knockdown models in PANC-1 and PATU 8988S cells exhibiting constitutively upregulated PKM expression (**Figure 2A**). Following rigorous screening, shPKM-1 and shPKM-3 constructs exhibiting optimal silencing efficiency were utilized for lentiviral transduction (**Figure S1A**). Stable PKM-depleted clones were validated via fluorescence-activated cell analysis of infection efficiency (>95% GFP+) (**Figure S1B**) coupled with qRT-PCR (**Figure 2B**) and immunoblot (**Figure 2C**) verification of PKM suppression.

**Figure 2.**
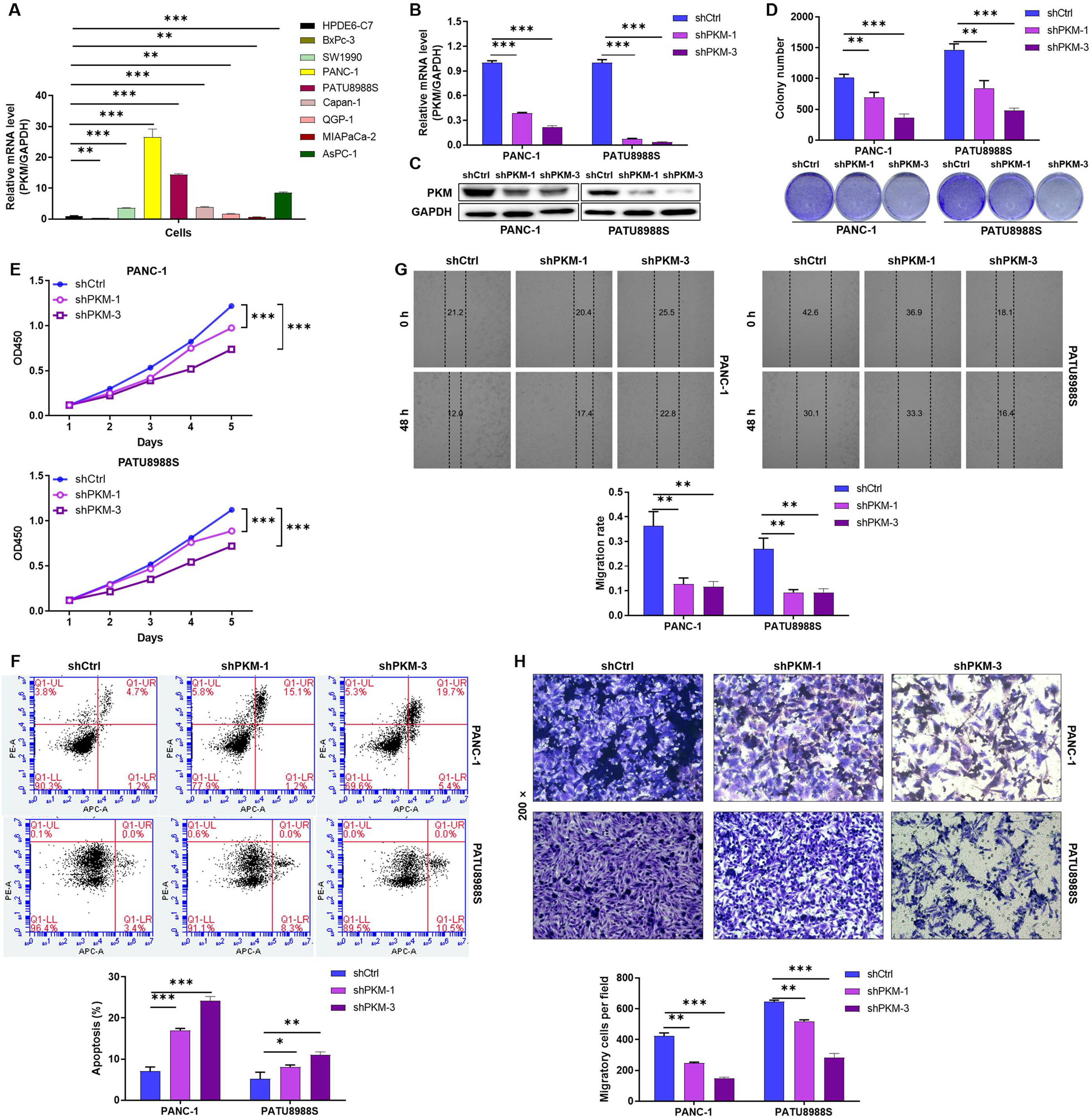
Knockdown of O-GlcNAcylation PKM inhibits the proliferation and migration of PDAC cells. **(A)** Relative mRNA expression levels of PKM in various pancreatic cancer cell lines (PANC-1, PATU8988S, SW1990, Capan-1, AsPC-1, MIA PaCa-2, and BxPC-3) compared to normal human pancreatic duct epithelial cells (HPDE6-C7). **(B)** Quantitative real-time PCR analysis confirming PKM mRNA knockdown efficiency in PANC-1 and PATU8988S cells transduced with shPKM-1, shPKM-3, or control shRNA (shCtrl). Results are normalized to GAPDH expression. **(C)** Western blot analysis of PKM protein expression in PANC-1 and PATU8988S cells with PKM knockdown (shPKM-1 and shPKM-3) compared to controls (shCtrl). GAPDH served as loading control. **(D)** Colony formation assay showing reduced clonogenic survival in PKM-knockdown PANC-1 and PATU8988S cells compared to controls. Colony numbers were quantified and presented as bar graph (right panel). **(E)** CCK-8 proliferation assays demonstrating significantly reduced cell viability in PKM-knockdown PANC-1 and PATU8988S cells compared to controls over 5 days of culture. **(F)** Flow cytometric analysis of apoptosis using Annexin V-FITC/PI staining in PKM-knockdown PANC-1 and PATU8988S cells compared to controls. The percentage of apoptotic cells (Q1-UR+Q1-LR) was quantified. **(G)** Wound healing assays showing reduced migratory capacity in PKM-knockdown PANC-1 and PATU8988S cells compared to controls at 48h post-scratch. Quantification of migrated cells is shown in the right panel. **(H)** Transwell Matrigel invasion assays illustrating decreased invasive potential of PKM-knockdown PANC-1 and PATU8988S cells compared to controls. Quantification of migrated cells is shown in the right panel. All experiments were performed in triplicate. Statistical significance was determined by two-tailed Student’s t-test. Data are presented as mean ± SD (n=3). *P<0.05, **P<0.01, ***P<0.001.

Subsequently, clonogenic survival assays demonstrated substantially attenuated colony-forming capacity (P<0.01) (**Figure 2D**). CCK-8 assays revealed that PKM ablation elicited significant antiproliferative effects (P<0.001), reducing viability in PANC-1 and PATU 8988S cells respectively over a 5-day observation period (**Figure 2E**). Furthermore, Annexin V/PI staining coupled with Flow cytometry analysis quantified a 2-fold increase in apoptotic rates (P<0.05), revealed that PKM knockdown promoted apoptosis in PDAC cells (**Figure 2F**). Functional interrogation of metastatic potential through wound healing assays showed suppressed migratory capacity (P<0.01) **(Figure 2G)** while Transwell Matrigel invasion assays revealed a reduction in invasive potential (P<0.001) **(Figure 2H)**. These data collectively establish that PKM ablation impedes PDAC progression across proliferation, survival, and metastatic axes, validating its oncogenic driver role in pancreatic carcinogenesis.

### O-GlcNAcylated PKM-Driven Metabolic Reprogramming Promotes PDAC Progression Through Aerobic Glycolysis

Mechanistic analysis revealed that PKM overexpression induces profound glycolytic flux remodeling in pancreatic ductal adenocarcinoma cells, characterized by: increase in glucose uptake (p<0.001; **Figure 3A**), elevation in ATP generation via substrate-level phosphorylation (p<0.001; **Figure 3B**),with lactate secretion increase (p<0.001; **Figure 3C**). Pharmacological inhibition using 2-deoxy-D-glucose (2-DG, 10 mM) effectively reversed PKM-mediated Warburg effect, reducing glycolytic intermediates accumulation in glucose uptake (p<0.001; **Figure 3A**), ATP generation (p<0.001; **Figure 3B**) with lactate secretion (p<0.001; **Figure 3C**). Functional rescue experiments confirmed metabolic dependency, with 2-DG treatment, reduced proliferation rates (p<0.001; **Figure 3D**), inhibited migration capacity (p<0.01; **Figure 3E**). Collectively, this study establishes that PKM serves as a metabolic gatekeeper in PDAC progression.

**Figure 3.**
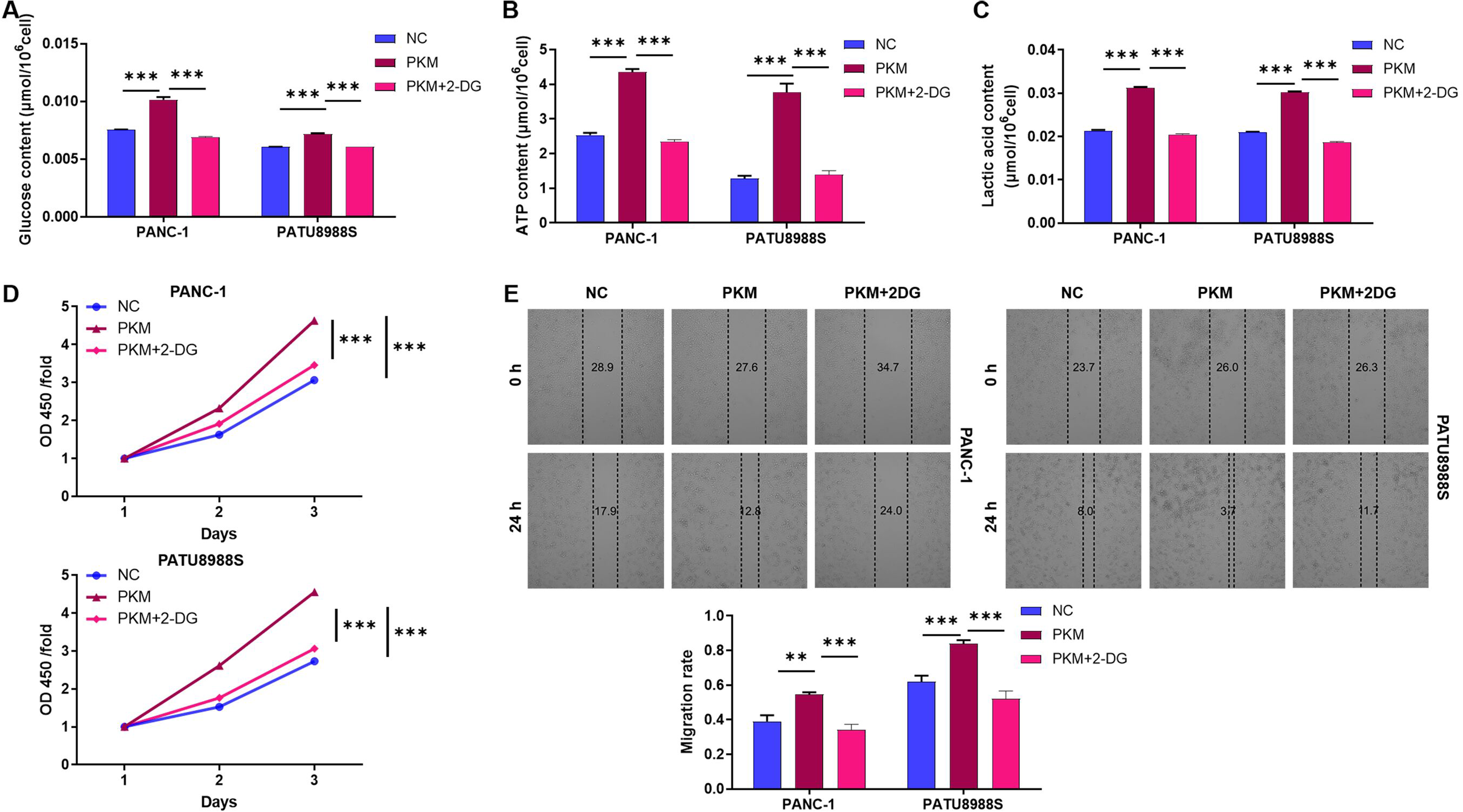
O-GlcNAcylation PKM overexpression induces metabolic reprogramming and promotes cell proliferation and migration in pancreatic cancer cells, which can be reversed by 2-DG treatment. **(A)** PKM overexpression significantly increased glucose uptake in both PANC-1 and PATU8988S cells compared to the negative control (NC) group. This effect was markedly reduced upon treatment with 2-deoxy-D-glucose (2-DG), a glycolysis inhibitor. **(B)** Elevation of ATP production through substrate-level phosphorylation was observed in PKM-overexpressing PANC-1 and PATU8988S cells. Treatment with 2-DG significantly attenuated PKM-induced ATP generation. **(C)** PKM overexpression substantially increased lactate secretion in both pancreatic cancer cell lines. This glycolytic phenotype was effectively reversed by 2-DG treatment. **(D)** PKM overexpression significantly promoted cell proliferation in both PANC-1 and PATU8988S cells, as evidenced by increased optical density (OD) values at 450 nm. The proliferative advantage induced by PKM was markedly suppressed by 2-DG treatment. **(E)** PKM overexpression significantly enhanced the migratory capacity of PANC-1 and PATU8988S cells. Treatment with 2-DG effectively reduced the migration rate of PKM-overexpressing cells. All experiments were performed in triplicate. Statistical significance was determined by two-tailed Student’s t-test. Data are presented as mean ± SD (n=3). *P<0.05, **P<0.01, ***P<0.001.

### Thr50 O-GlcNAcylation Governs PKM-Driven Metabolic Plasticity and Oncogenesis in PDAC Cells

Structural bioinformatics analysis (NetOGlyc 4.0) predicted three conserved O-GlcNAcylation motifs in PKM: Thr50 (PPV48T50AT), Thr405 (T405VSS), and Ser406 (VS406SV). Site-directed mutagenesis generated four Flag-PKM variants in pcDNA3.1: WT (T50/T405/S406), T50A (Thr50→Ala), T405A (Thr405→Ala) and S406A (Ser406→Ala). After transfecting these constructs into HEK293T cells, Co-IP with anti-Flag antibody showed the reduction in O-GlcNAc signal for T405A/S406A mutants, loss of O-GlcNAc signal in T50A mutant (**Figure 4A**). This identifies Thr50 as the dominant O-GlcNAc acceptor site in PKM.

**Figure 4.**
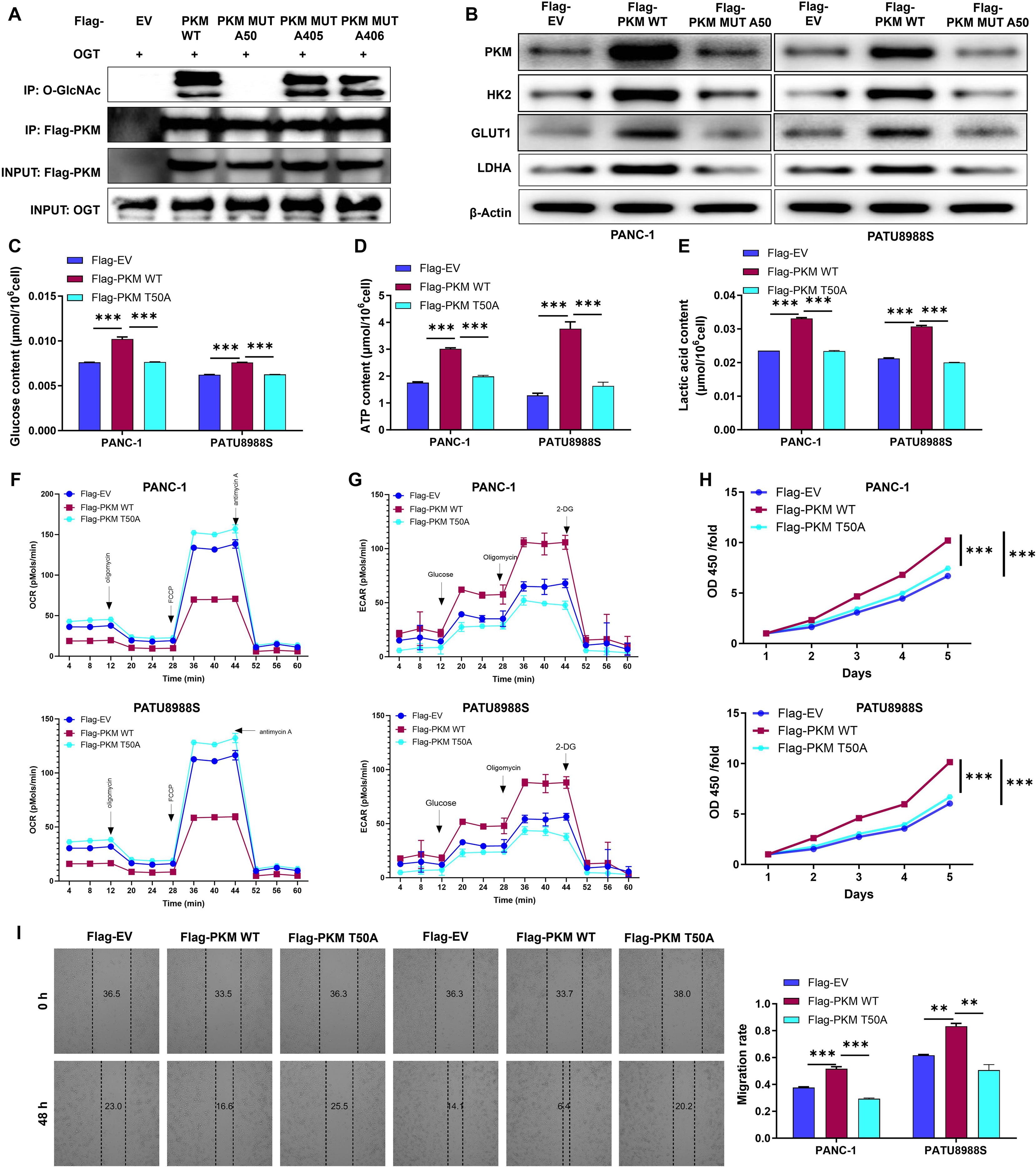
Thr50 O-GlcNAcylation Regulates PKM2-Mediated Metabolic Reprogramming and Oncogenic Properties in PDAC Cells. **(A)** Co-immunoprecipitation (Co-IP) analysis showing O-GlcNAcylation levels of Flag-PKM wild-type (WT) and mutant variants (T50A, T405A, S406A) in HEK293T cells. IP was performed using anti-Flag antibody followed by O-GlcNAc Western blotting (upper panel). Total PKM protein levels were determined by Western blotting (middle panel), and OGT expression in input samples was confirmed (lower panel). **(B)** Western blot analysis of key glycolytic enzymes (HK2, GLUT1, and LDHA) in PANC-1 cells expressing Flag-PKM WT or mutants. β-Actin served as loading control. **(C-G)** Quantitative analysis of metabolic parameters in PANC-1 and PATU8988S cells expressing PKM variants: **(C)** Glucose uptake, **(D)** ATP content, **(E)** Lactate secretion, **(F)** Extracellular acidification rate (ECAR), and **(G)** Oxygen consumption rate (OCR). Statistical significance was determined by two-tailed Student’s t-test. **(H-I)** Proliferation (**H)** and migration **(I)** assays of PANC-1 and PATU8988S cells stably expressing PKM WT or T50A mutant. Data presented as mean ± SD (n = 3 independent experiments). *P<0.05, **P<0.01, ***P<0.001.

Immunoblot analysis revealed PKM2 O-GlcNAcylation upregulated key metabolic enzyme expression, Hexokinase 2 (HK2), Glucose transporter 1 (GLUT1), Lactate dehydrogenase A (LDHA) **(Figure 4B**). Furthermore, metabolic phenotyping in PANC-1 and PATU 8988S cells demonstrated that WT PKM2 overexpression induced Warburg effect remodeling: glucose uptake increased (p<0.001; **Figure 4C**), ATP levels elevated (p<0.001; **Figure 4D**), lactate secretion increased (p<0.001; **Figure 4E**), ECAR increased (**Figure 4F, S2A-S2C**), OCR decreased (**Figure 4G, S2D-S2E**). Notably, following T50A mutation, the effect is no longer observable **(Figure 4B-4G**). These findings demonstrate that Thr50 O-GlcNAcylation is essential for PKM-mediated metabolic reprogramming in PDAC cells.

Consistent with the metabolic findings, functional assays showed T50A mutant failed to stimulate proliferation (p<0.001; **Figure 4H**) and promote migration (p<0.01; **Figure 4I**) in PDAC cells, further confirming the essential role of Thr50 modification in PKM-mediated oncogenic progression. The integrated results demonstrate that O-GlcNAcylation at the Thr50 site of PKM plays a decisive role and is essential for regulating glycolytic activity and malignant biological behaviors in PDAC cells.

### Thr50 O-GlcNAcylation-Dependent Regulation of CDC27 Transcription via ARNT by PKM in PDAC

To elucidate the downstream mechanisms by which O-GlcNAcylated PKM drives pancreatic carcinogenesis, we performed tandem mass tag (TMT)-based quantitative proteomics in PKM-knockdown PDAC cells, identifying CDC27 as a top downregulated gene alongside Tax1bp1, Pcm1, Rnf40, and Chd1. Correlation analysis of TCGA PDAC cohorts revealed a robust positive correlation between PKM and CDC27 mRNA (Pearson’s r = 0.26, p < 0.001; **Figure 5A**) and protein levels **(Figure 5B)**. To determine whether O-GlcNAcylation regulates PKM-mediated CDC27 regulation, we ectopically expressed wild-type (WT) or Thr50 O-GlcNAcylation-deficient (Mut A50) PKM in PDAC cells. While WT PKM overexpression robustly upregulated CDC27 mRNA and protein **(Figure 5C)**, the Mut A50 variant completely abolished this effect, establishing O-GlcNAcylation at Thr50 as critical for PKM-driven CDC27 activation.

**Figure 5.**
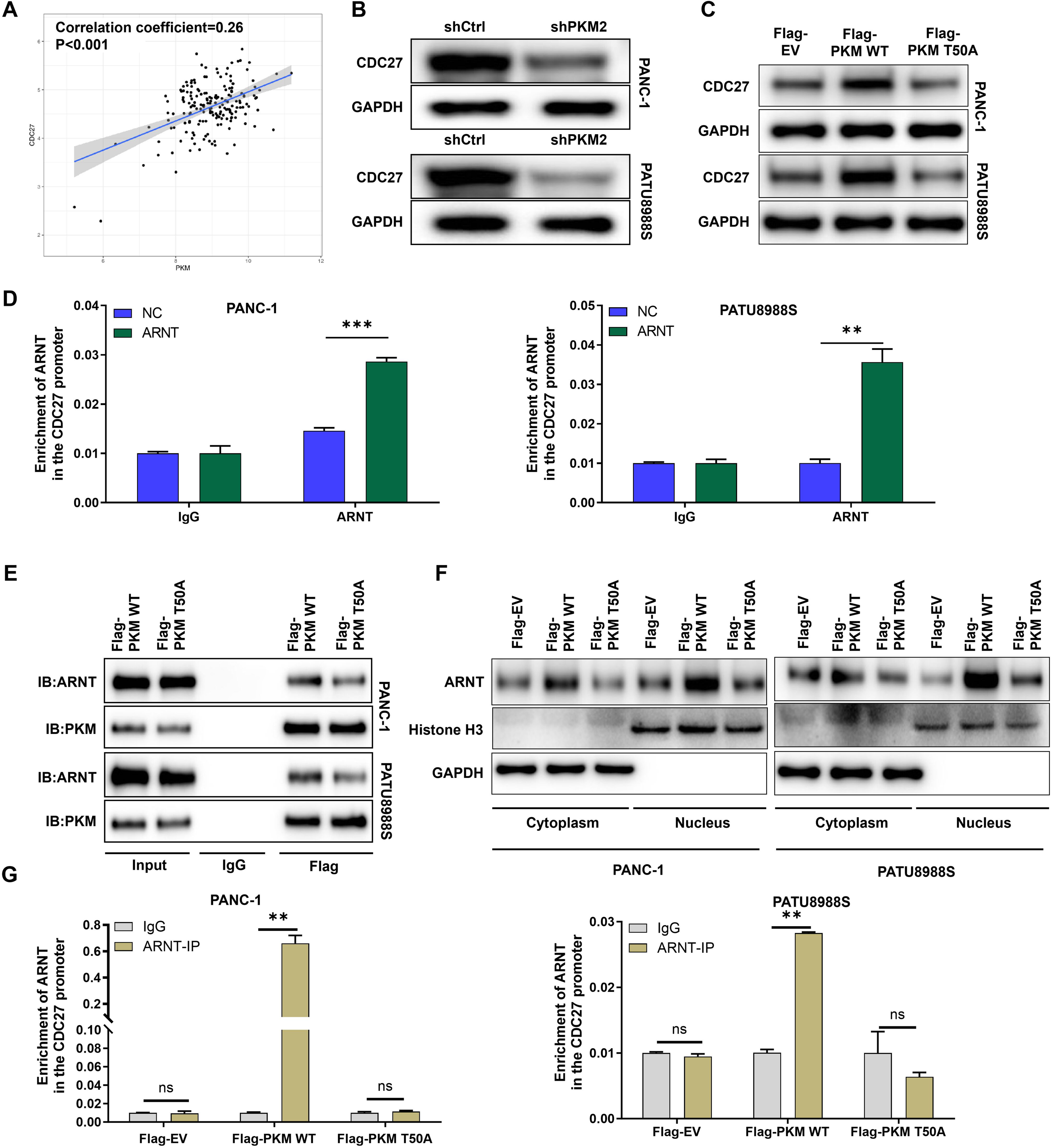
Thr50 O-GlcNAcylation-Dependent Regulation of CDC27 Transcription via ARNT by PKM in PDAC. **(A)** Scatter plot illustrating the positive correlation between PKM and CDC27 mRNA expression in TCGA PDAC cohorts (Pearson’s r = 0.26, p < 0.001). **(B)** Western blot analysis showing decreased CDC27 expression in shPKM2-transfected PDAC cells compared to shCtrl controls. **(C)** Western blot analysis demonstrates that overexpression of wild-type PKM (PKM WT) significantly upregulates CDC27 expression, whereas the O-GlcNAcylation-deficient mutant (PKM T50A) fails to induce CDC27 expression. GAPDH served as a loading control. **(D)** ChIP-qPCR results showing ARNT enrichment at the CDC27 promoter region in PANC-1 and PATU8988S cells. (**E**) Co-IP analysis showing the interaction between ARNT and PKM in PANC-1 and PATU8988S cells with Flag-PKM WT and PKM T50A. **(F)** Nuclear-cytoplasmic fractionation followed by western blot analysis showing that PKM WT, but not PKM T50A, promotes ARNT nuclear translocation. Histone H3 and GAPDH were used as nuclear and cytoplasmic markers, respectively. **(G)** Quantitative analysis of ChIP-qPCR data confirming that ARNT occupancy at the CDC27 promoter was significantly reduced in cells expressing PKM T50A compared to PKM WT. Data are presented as mean ± SD from three independent experiments. ns: no significance, **P<0.01, ***P<0.001.

To explore the mechanism by which PKM regulates CDC27, using intersecting interactome (STRING database) and transcription factor prediction (hTFtarget database) analyses, ARNT emerged as a candidate mediator. Chromatin immunoprecipitation (ChIP)-qPCR demonstrated ARNT binding to the CDC27 promoter region **(Figure 5D)**. Co-IP revealed a physical association between PKM and ARNT, which was disrupted by Thr50 O-GlcNAcylation mutation **(Figure 5E)**. Notably, nuclear fractionation revealed that PKM WT, but not Mut A50, promoted ARNT nuclear translocation **(Figure 5F)**, suggesting O-GlcNAcylation regulates PKM’s ability to recruit ARNT into the nucleus. Functional validation through ChIP-qPCR in PKM WT/Mut A50-overexpressing cells confirmed that ARNT enrichment at the CDC27 promoter was abolished by Thr50 O-GlcNAcylation mutation **(Figure 5G)**. Collectively, these findings reveal an O-GlcNAcylation-dependent axis where PKM modulates ARNT-mediated CDC27 transcription, thereby driving PDAC progression.

### PKM-Driven AKT Activation via CDC27 Facilitates Aerobic Glycolysis and Progression in PDAC

IHC staining of clinical specimens further demonstrated elevated CDC27 protein expression in pancreatic tumors compared to adjacent normal tissues **(Figure 6A)**, with high CDC27 levels correlating with poor overall survival (p = 0.015; **Figure 6B**). To clarify that PKM drives the progression of PDAC through CDC27, PKM overexpression and CDC27 knockdown were constructed respectively in the designated cells (**Figure S3A-S3C**). Phenotypic rescue experiments validated this dependency. PKM overexpressing cells exhibited enhanced proliferation (p<0.05; **Figure 6C**) and migration (p<0.001; **Figure 6D**). Furthermore, CDC27 knockdown reduced proliferation (p<0.05;) and migration (p<0.05) of PKM overexpressing (**Figure 6C-6D**). Moreover, CDC27 knockdown downregulated key glycolytic enzymes (GLUT1, HK2, LDHA) induced by PKM overexpressing **(Figure 6E)**. Meanwhile, CDC27 knockdown partially rescued PKM overexpressing induced glycolytic activation, such as glucose uptake (p<0.001; **Figure 6F**), ATP production (p<0.01; **Figure 6G**), and lactate secretion (p<0.001; **Figure 6H**). Similarly, ECAR and OCR were normalized to control levels upon CDC27 suppression (**Figure 6I–6J, S4A-S4F**). These data demonstrated that PKM’s oncogenic effects are mediated through CDC27.

**Figure 6.**
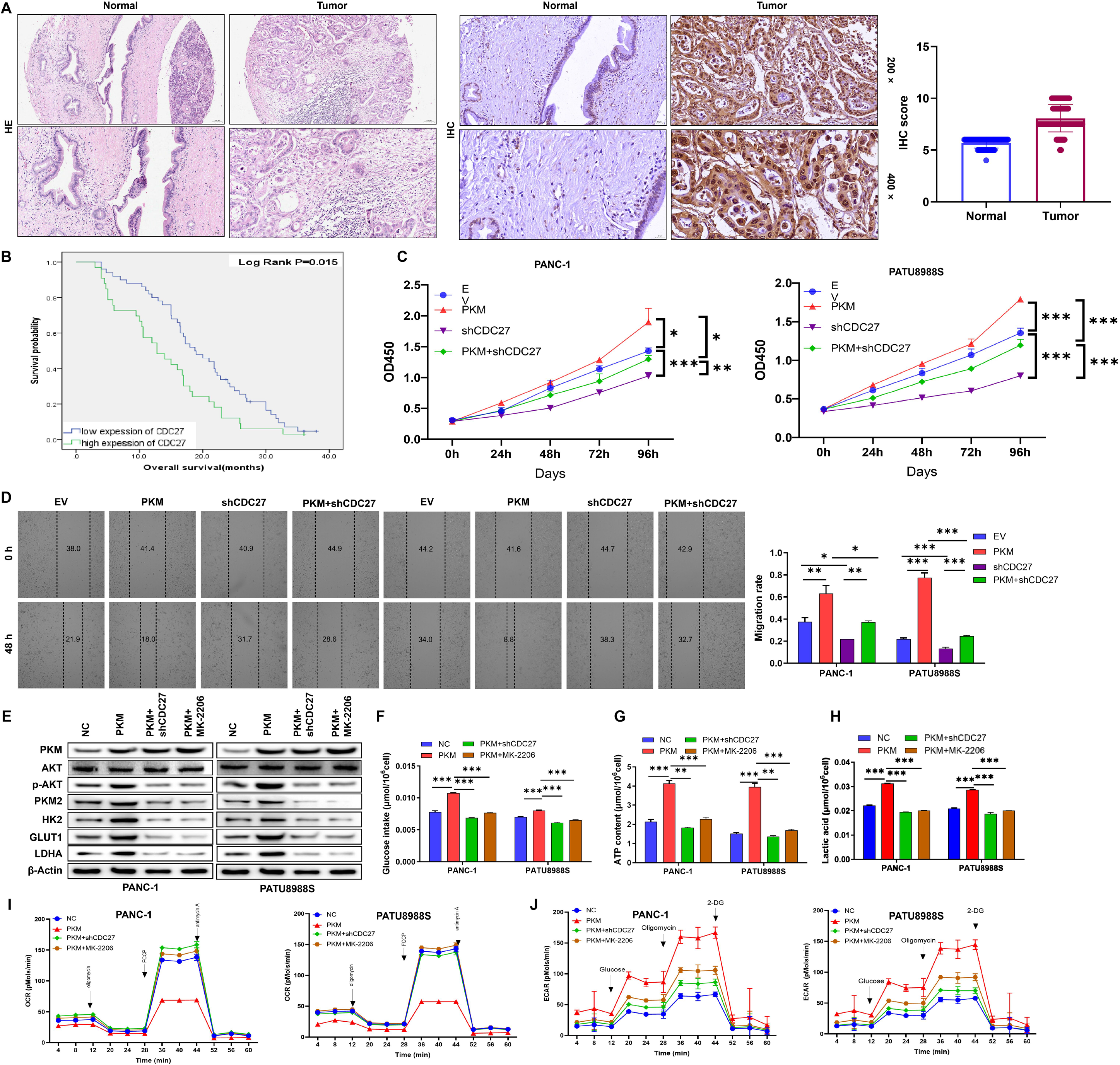
PKM-driven AKT activation via CDC27 promotes PDAC progression and aerobic glycolysis. **(A)** Hematoxylin-eosin (HE) staining and immunohistochemistry (IHC) analysis of CDC27 expression in paired normal pancreatic tissues and pancreatic ductal adenocarcinoma (PDAC) specimens. Scale bar: 100 μm. Right panel: Quantification of CDC27 IHC scores in normal and tumor tissues. **(B)** Kaplan-Meier overall survival curves for PDAC patients with low versus high CDC27 expression. Log-rank test, P = 0.015. **(C)** Proliferation curves of PANC-1 and PATU8988S cells stably transfected with empty vector (EV), PKM overexpression (PKM), CDC27 knockdown (shCDC27), or both PKM overexpression and CDC27 knockdown (PKM+shCDC27). **(D)** Wound healing assays demonstrating migration ability of PANC-1 and PATU8988S cells under indicated conditions. Quantification of migrated cells is shown on the right. Scale bar: 200 μm. **(E)** Western blot analysis of p-AKT, AKT, PKM2, GLUT1, HK2, LDHA in cells with indicated treatments. Both PKM overexpression and AKT inhibited (PKM+MK2206) settled as the fourth group, β-actin served as loading control. **(F-J)** Analysis of glucose metabolism in PANC-1 and PATU8988S cells with indicated treatments: **(F)** glucose uptake, **(G)** ATP production, **(H)** lactate secretion, **(I)** extracellular acidification rate (ECAR), and **(J)** oxygen consumption rate (OCR). Data are presented as mean ± SD from three independent experiments. *P<0.05, **P<0.01, ***P<0.001.

Given the established oncogenic role of AKT in metabolic reprogramming, we investigated whether CDC27 modulates AKT signaling in PDAC. In the study, knockdown of CDC27 in PDAC cells can counteract the level of phosphorylated AKT (p-AKT) induced by PKM overexpression (**Figure S3D**), suggested that PKM drives AKT activation through CDC27. Furthermore, the addition of AKT pathway inhibitors (MK-2206) effectively mitigates the glycolytic activity alterations induced by PKM overexpression. This is evidenced by the downregulation of key enzyme expression levels **(Figure 6E)**, reduced glucose uptake (p<0.001; **Figure 6F**), ATP production (p<0.001; **Figure 6G**), and lactic acid secretion (p<0.001; **Figure 6H**), increased OCR (**Figure 6I**), and decreased ECAR (**Figure 6J**). Collectively, this study elucidates a novel mechanism where PKM facilitates PDAC progression by CDC27, which in turn sustains AKT signaling to promote glycolytic addiction and malignant phenotypes.

### CDC27 knockdown upregulates PPP2CA by reducing its K11-linked ubiquitination and decreases AKT phosphorylation

First, we confirmed that CDC27 interacts with PPP2CA in two distinct pancreatic cancer cell lines (PANC-1 and PATU 8988S)via co-immunoprecipitation (Co-IP)**(Fig7A)**. Knockdown CDC27 significantly upregulated PPP2CA expression**(Fig7B)**, while markedly reducing the K11-linked ubiquitination of PPP2CA**(Fig7C)**. Furthermore, knockdown of PPP2CA significantly increased the phosphorylation level of AKT**(Fig7D)**. These results suggest that reduced CDC27 expression may upregulate PPP2CA expression by decreasing K11-linked ubiquitination of PPP2CA, thereby lowering AKT phosphorylation. This indicates that PPP2CA may be involved in the negative regulation of the AKT signaling pathway.

### Inhibiting Thr50 O-GlcNAcylation of PKM significantly reduced the proliferation of pancreatic tumor in vivo

We observed that tumor diameter, volume, and weight in the PKM wild-type (WT) group were significantly higher than those in the control group and the O-GlcNAcylation-inhibited group**(Fig 8A-8C)**, indicating that O-GlcNAcylated PKM promotes pancreatic tumor proliferation. No significant difference in tumor proliferation was observed between the PKM O-GlcNAcylation-inhibited group and the empty vector group. Immunohistochemical staining revealed that the expression of CDC27 and Ki67 were significantly increased in the PKM WT group, while PPP2CA decreased**(Fig 8D)**. After inhibiting PKM O-GlcNAcylation, the expression of PKM showed no obvious change compared with the WT group, whereas the expression of CDC27 and Ki67 was obviously deceased, which were close to that in the empty vector group, while PPP2CA presented the opposite result. Further Western blot (WB) analysis indicated that the expression levels of CDC27, phosphorylated AKT (p-AKT), and HK2 were significantly elevated in tumor tissues of the PKM WT group. In contrast, inhibition of Thr50 O-GlcNAcylation PKM resulted in the opposite effect, with markedly decreased expression of CDC27, p-AKT, and HK2**(Fig 8E)**.

## Discussion

Our study identifies O-GlcNAcylated PKM as a pivotal metabolic driver in pancreatic ductal adenocarcinoma. By integrating proteomic, transcriptomic, and clinical analyses, we demonstrate that PKM is significantly upregulated in pancreatic tumors compared to normal tissues (Figures 1B–1I), correlating with advanced tumor stage, metastasis, and poor prognosis. This aligns with prior studies highlighting PKM’s role in metabolic reprogramming and tumorigenesis across cancers. However, our work advances this understanding by elucidating the critical role of O-GlcNAcylation at Thr50 in PKM’s oncogenic activity. PKM produces two completely independent protein monomers through mutually exclusive alternative splicing: retention of exon 9 translates into the PKM1 monomer, whereas retention of exon 10 translates into the PKM2 monomer. Bioinformatic analysis predicts that the glycosylation sites of PKM include Thr45 and Thr50 (located in exon 2 at the N-terminus of PKM2), as well as Thr405, Ser406 and Tyr406 (located in exon 10 at the C-terminus of PKM2). Unlike previous reports focusing on PKM’s canonical metabolic functions(13–15), and O-GlcNAc transferase (OGT)-mediated post-translational modification of PKM2 at Thr405, Ser406 and Tyr406 stabilizes its dimeric conformation, we reveal that O-GlcNAcylation at Thr50 site modulates PKM2’s transcriptional and interatomic landscape, enabling its crosstalk with oncogenic pathways such as AKT and CDC27. We identify CDC27, a key regulator of mitotic exit and genomic stability, as a novel transcriptional target of PKM2. While CDC27 has been implicated in cancer progression through its role in cell cycle regulation(10, 16, 17), our findings establish a unique mechanism where Thr50 O-GlcNAcylated PKM2 drives CDC27 expression via ARNT-mediated transcriptional activation (Figures 5A–5G). This contrasts with studies linking CDC27 to tumor suppression in certain contexts(18, 19), underscoring the context-dependent nature of its function. The dependence of CDC27 upregulation on PKM2’s Thr50 O-GlcNAcylation (Figure 5B) suggests a paradigm shift in how post-translational modifications rewire transcriptional networks in cancer.

The discovery of ARNT as a mediator of PKM2-CDC27 signaling introduces a novel axis in PDAC biology. ARNT, a transcription factor central to hypoxia adaptation and metabolic regulation(20, 21), has not been previously linked to PKM2 or CDC27. Our ChIP and co-IP analyses demonstrate that O-GlcNAcylated PKM2 facilitates ARNT nuclear translocation and promoter binding (Figures 5C–5E), paralleling studies where ARNT drives oncogenic gene expression in hypoxic tumors(22–25). However, our work uniquely positions ARNT as a bridge between PKM2’s metabolic activity and cell cycle regulation, revealing a multi-functional role in PDAC progression.

By elucidating the T50A-PKM2-CDC27-PPP2CA-AKT cascade, we provide mechanistic insight into how O-GlcNAcylation sustains glycolytic addiction in PDAC. Our results suggest that inhibiting O-glycosylation fo PKM2 at the Thr50 site reduces CDC27 expression, which leads to downregulation of ubiquitination at the K11 site of PPP2CA, resulting in upregulation of PPP2CA expression (Figures 7A–7D). This, in turn, reduces AKT phosphorylation, ultimately inhibiting glucose metabolism and pancreatic tumor proliferation in vivo (Figures 8A–8E). CDC27 knockdown attenuates AKT phosphorylation and glycolytic flux (Figures 6A–6G), mirroring observations in cancers where AKT hyperactivation drives metabolic plasticity(26–28). Notably, the rescue of proliferative and migratory phenotypes by WT PKM—but not Thr50-mutant—highlights the indispensability of O-GlcNAcylation in maintaining this axis (Figure 6H–6I). These findings are also confirmed in vivo (Figures 8A–8E). This aligns with emerging evidence that O-GlcNAcylation regulates oncogenic kinase signaling(29–31), yet expands its functional repertoire to include transcriptional crosstalk with metabolic enzymes.

**Figure 7.**
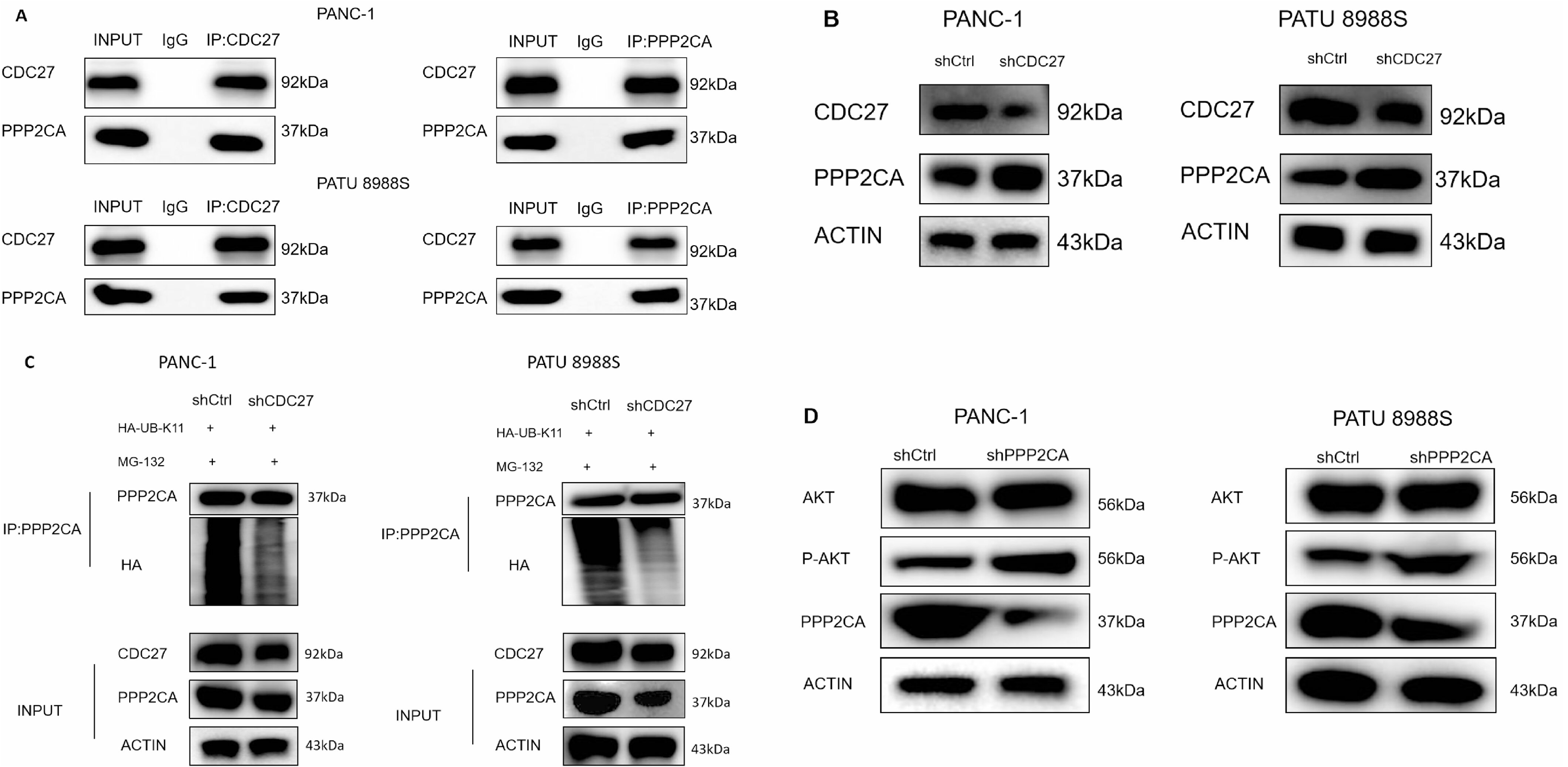
CDC27 knockdown upregulates PPP2CA by reducing its K11-linked ubiquitination and decreases AKT phosphorylation. (A)CDC27 interacts with PPP2CA in two distinct pancreatic cancer cell lines (PANC-1 and PATU 8988S)via co-immunoprecipitation (Co-IP). (**B**)Knockdown CDC27 significantly upregulated PPP2CA expression, (**C**)Knockdown CDC27 markedly reducing the K11-linked ubiquitination of PPP2CA. (**D**)knockdown PPP2CA significantly increased the phosphorylation level of AKT.

**Figure 8.**
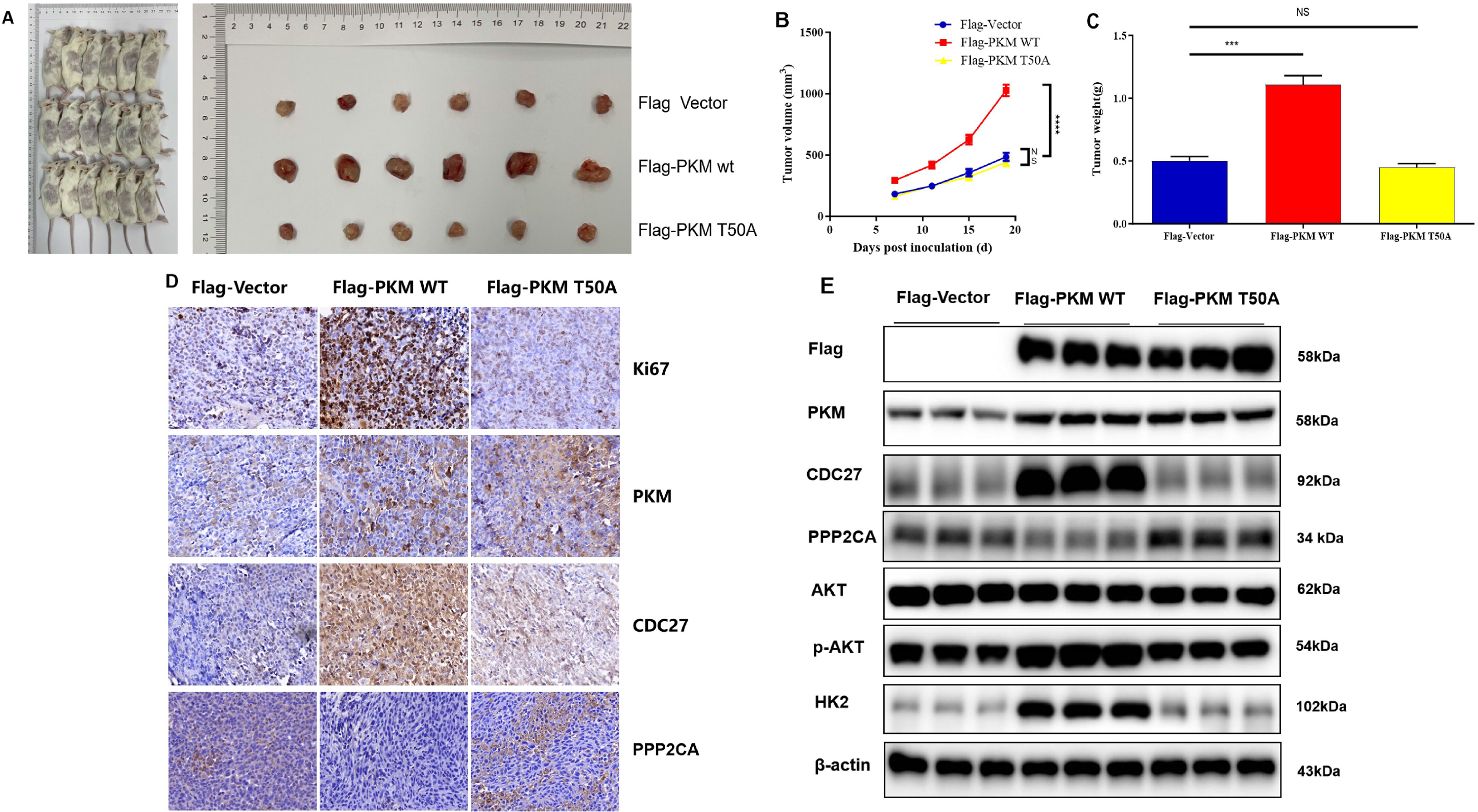
Inhibiting Thr50 O-GlcNAcylation of PKM significantly reduced the proliferation of pancreatic tumor in vivo. (A,B,C) Tumor diameter, volume, and weight in the PKM wild-type (WT) group were significantly higher than those in the control group and the O-GlcNAcylation-inhibited group. No significant difference in tumor proliferation was observed between the PKM O-GlcNAcylation-inhibited group and the empty vector control group. **(D)** Immunohistochemical staining revealed that the expression of CDC27 and Ki67 were significantly increased in the PKM WT group, while PPP2CA decreased. After inhibiting PKM O-GlcNAcylation, the expression of PKM2 showed no obvious change compared with the WT group, whereas the expression of CDC27 and Ki67 was obviously deceased, which were close to that in the empty vector control group, while PPP2CA presented the opposite result. **(E)** Western blot (WB) analysis indicated that the expression levels of CDC27, phosphorylated AKT (p-AKT), and HK2 were significantly elevated in tumor tissues of the PKM WT group. In contrast, inhibition of PKM O-GlcNAcylation resulted in the opposite effect, with markedly decreased expression of CDC27, p-AKT, and HK2.

Clinically, our data position O-GlcNAcylated PKM as a dual prognostic biomarker and therapeutic target. The correlation between PKM expression and adverse outcomes (Figure 1E–1I) mirrors findings in breast and lung cancers(32–34), reinforcing its broad relevance in malignancy. Targeting the Thr50-PKM2-CDC27-PPP2CA-AKT axis could disrupt the metabolic vulnerability of PDAC, particularly given the limited efficacy of current therapies against this aggressive cancer. However, challenges remain, including potential off-target effects on ARNT-dependent hypoxia responses and the need to validate these findings in patient-derived models.

While our study advances the mechanistic understanding of PKM in PDAC, several limitations warrant mention. First, the clinical cohort analyzed is relatively small; larger validation cohorts are needed to confirm prognostic significance. Second, the in vitro and in vivo models used may not fully recapitulate the tumor microenvironment’s complexity. Experiments in vivo, shows inhibiting the Thr50 O-GlcNAcylation of PKM2 can induce expression regulation of downstream proteins such as CDC27 and AKT, as well as functional alterations in glucose metabolism and tumor proliferation, while has little effect on PKM2 expression(Figure 8D, E). This may be related to that the Thr50 site is located in exon2 of PKM2(In contrast, Thr405, Ser406, and Tyr406 are located in exon10, and inhibition of these sites leads to reduced PKM2 expression). Future work should verify this and explore how Thr50 O-GlcNAcylation mutations affect PKM2’s interaction with other metabolic enzymes could deepen insights into metabolic reprogramming.

In summary, we delineate a novel O-GlcNAcylation-dependent pathway where PKM drives PDAC progression through CDC27 transcriptional activation and AKT-mediated glycolysis. This work not only elucidates molecular mechanisms underlying metabolic oncogenesis but also identifies actionable targets for therapeutic intervention in PDAC.

## Author contributions

Bin Yang, Yi Zhu and Xiaopeng Zhan contributed equally to this work. Shikai Zhu(the primary corresponding author) and Yu Zhou are co-corresponding authors. Bing Yang conceived the study, performed experiments, analyzed data, drafted and revised the manuscript. Yi Zhu participated in study design, experimental implementation and data analysis. Xiaopeng Zhan, Yun Zhang and Jianing Cui conducted experiments and analyzed data. Yu Zhou and Shikai Zhu contributed to study design, data analysis and manuscript revision. All the authors have confirmed the submission of this manuscript.

## Data availability statement

The data that support the findings of this study are available from the corresponding author upon reasonable request.

---

**Figure S1. PKM knockdown efficiency in PDAC cells. (A)** Relative mRNA levels of PKM in PANC-1 and PATU8988S cells stably expressing either control (CON, shCtrl) or PKM-targeting shRNAs (shPKM-1, shPKM-2, shPKM-3) were determined by qRT-PCR. Results are presented as mean ± SD from three independent experiments. **P<0.01, ***P<0.001. **(B)** Phase-contrast and fluorescence microscopy images showing cellular morphology and GFP expression (indicative of successful lentiviral transduction) in PANC-1 (left panel) and PATU8988S (right panel) cells stably expressing control shRNA (shCtrl) or two effective PKM-targeting shRNAs (shPKM-1 and shPKM-3).

**Figure S2. Thr50 O-GlcNAcylation of PKM2 regulates metabolic reprogramming in PDAC.** PANC-1 and PATU8988S cells stably expressing Flag-empty vector (Flag-EV), wild-type PKM2 (Flag-PKM WT), or mutant PKM2 with threonine 50 alanine substitution (Flag-PKM MUT (A50)) were analyzed for metabolic phenotypes. Bar graphs depict quantitative differences in glycolytic **(A-F)** and mitochondrial respiration **(G-I)** parameters. Results are presented as mean ± SD from three independent experiments. **P<0.01, ***P<0.001.

**Figure S3. PKM drives PDAC progression through CDC27-mediated AKT activation. (A-B)** Quantitative real-time PCR analysis of PKM, CDC27, and GAPDH mRNA expression levels in PANC-1 and PATU8988S cells with indicated treatments (EV, PKM overexpression, shCDC27 knockdown, and PKM+shCDC27 combination). Results are presented as mean ± SD from three independent experiments. ns: no significance, ***P<0.001. **(C-D)** Western blot analysis of PKM, CDC27, GAPDH, AKT, p-AKT, and β-actin protein expression levels in PANC-1 and PATU8988S cells with indicated treatments. β-actin served as a loading control.

**Figure S4. Effect of PKM overexpression, CDC27 knockdown and AKT inhibition on mitochondrial respiration and glycolysis in PDAC. (A-C)** Mitochondrial respiration parameters: (A) Basal respiration; (B) ATP-linked respiration; (C) Maximal respiration. **(D-F)** Glycolytic parameters: (D) Glycolysis; (E) Glycolysis capacity; (F) Glycolytic reserve. (EV, empty vector; PKM, PKM overexpression; shCDC27, CDC27 knockdown; and PKM+shCDC27, both PKM overexpression and CDC27 knockdown;. or PKM+MK2206, both PKM overexpression and AKT inhibited). Data are presented as mean ± SD from three independent experiments. *P<0.05, **P<0.01, ***P<0.001.

## Reference

1. Deplanque G, Demartines N. Pancreatic cancer: are more chemotherapy and surgery needed? Lancet. 2017;389(10073):985–6.

2. Luo W, Wang J, Chen H, Ye L, Qiu J, Liu Y, et al. Epidemiology of pancreatic cancer: New version, new vision. Chin J Cancer Res. 2023;35(5):438–50.

3. Ying H, Kimmelman AC, Bardeesy N, Kalluri R, Maitra A, DePinho RA. Genetics and biology of pancreatic ductal adenocarcinoma. Genes Dev. 2025;39(1-2):36–63.

4. Schwartz L, Supuran CT, Alfarouk KO. The Warburg Effect and the Hallmarks of Cancer. Anticancer Agents Med Chem. 2017;17(2):164–70.

5. Rihan M, Nalla LV, Dharavath A, Shard A, Kalia K, Khairnar A. Pyruvate Kinase M2: a Metabolic Bug in Re-Wiring the Tumor Microenvironment. Cancer Microenviron. 2019;12(2-3):149–67.

6. Wang Y, Liu J, Jin X, Zhang D, Li D, Hao F, et al. O-GlcNAcylation destabilizes the active tetrameric PKM2 to promote the Warburg effect. Proc Natl Acad Sci U S A. 2017;114(52):13732–7.

7. Bond MR, Hanover JA. A little sugar goes a long way: the cell biology of O-GlcNAc. J Cell Biol. 2015;208(7):869–80.

8. Zhu Y, Hart GW. Targeting O-GlcNAcylation to develop novel therapeutics. Mol Aspects Med. 2021;79:100885.

9. Mathew M, Nguyen NT, Bhutia YD, Sivaprakasam S, Ganapathy V. Correction: Mathew, et al. Metabolic Signature of Warburg Effect in Cancer: An Effective and Obligatory Interplay between Nutrient Transporters and Catabolic/Anabolic Pathways to Promote Tumor Growth. Cancers 2024, 16, 504. Cancers (Basel). 2024;16(9).

10. Kazemi-Sefat GE, Keramatipour M, Talebi S, Kavousi K, Sajed R, Kazemi-Sefat NA, et al. The importance of CDC27 in cancer: molecular pathology and clinical aspects. Cancer Cell Int. 2021;21(1):160.

11. Qiu L, Tan X, Lin J, Liu RY, Chen S, Geng R, et al. CDC27 Induces Metastasis and Invasion in Colorectal Cancer via the Promotion of Epithelial-To-Mesenchymal Transition. J Cancer. 2017;8(13):2626–35.

12. Feng Z, Zhang L, Zhou J, Zhou S, Li L, Guo X, et al. mir-218-2 promotes glioblastomas growth, invasion and drug resistance by targeting CDC27. Oncotarget. 2017;8(4):6304–18.

13. Traxler L, Herdy JR, Stefanoni D, Eichhorner S, Pelucchi S, Szücs A, et al. Warburg-like metabolic transformation underlies neuronal degeneration in sporadic Alzheimer’s disease. Cell Metab. 2022;34(9):1248–63.e6.

14. Ma WK, Voss DM, Scharner J, Costa ASH, Lin KT, Jeon HY, et al. ASO-Based PKM Splice-Switching Therapy Inhibits Hepatocellular Carcinoma Growth. Cancer Res. 2022;82(5):900–15.

15. Bian Z, Yang F, Xu P, Gao G, Yang C, Cao Y, et al. LINC01852 inhibits the tumorigenesis and chemoresistance in colorectal cancer by suppressing SRSF5-mediated alternative splicing of PKM. Mol Cancer. 2024;23(1):23.

16. Lee SJ, Langhans SA. Anaphase-promoting complex/cyclosome protein Cdc27 is a target for curcumin-induced cell cycle arrest and apoptosis. BMC Cancer. 2012;12:44.

17. Xin Y, Ning S, Zhang L, Cui M. CDC27 Facilitates Gastric Cancer Cell Proliferation, Invasion and Metastasis via Twist-Induced Epithelial-Mesenchymal Transition. Cell Physiol Biochem. 2018;50(2):501–11.

18. Wang YX, Huang CY, Chiu HJ, Huang PH, Chien HT, Jwo SH, et al. Nuclear-localized CTEN is a novel transcriptional regulator and promotes cancer cell migration through its downstream target CDC27. J Physiol Biochem. 2023;79(1):163–74.

19. Zhu J, Liu X, Luan Z, Xue W, Cui H, Zhang B, et al. Circular RNA circSLC8A1 inhibits the proliferation and invasion of glioma cells through targeting the miR-214-5p/CDC27 axis. Metab Brain Dis. 2022;37(4):1015–23.

20. Mandl M, Depping R. Hypoxia-inducible aryl hydrocarbon receptor nuclear translocator (ARNT) (HIF-1â): is it a rare exception? Mol Med. 2014;20(1):215–20.

21. Wolff M, Jelkmann W, Dunst J, Depping R. The Aryl Hydrocarbon Receptor Nuclear Translocator (ARNT/HIF-1â) is influenced by hypoxia and hypoxia-mimetics. Cell Physiol Biochem. 2013;32(4):849–58.

22. Lee SH, Golinska M, Griffiths JR. HIF-1-Independent Mechanisms Regulating Metabolic Adaptation in Hypoxic Cancer Cells. Cells. 2021;10(9).

23. Alafate W, Lv G, Zheng J, Cai H, Wu W, Yang Y, et al. Targeting ARNT attenuates chemoresistance through destabilizing p38α-MAPK signaling in glioblastoma. Cell Death Dis. 2024;15(5):366.

24. Huang CR, Lee CT, Chang KY, Chang WC, Liu YW, Lee JC, et al. Down-regulation of ARNT promotes cancer metastasis by activating the fibronectin/integrin β1/FAK axis. Oncotarget. 2015;6(13):11530–46.

25. Toledo RA, Jimenez C, Armaiz-Pena G, Arenillas C, Capdevila J, Dahia PLM. Hypoxia-Inducible Factor 2 Alpha (HIF2á) Inhibitors: Targeting Genetically Driven Tumor Hypoxia. Endocr Rev. 2023;44(2):312–22.

26. Hoxhaj G, Manning BD. The PI3K-AKT network at the interface of oncogenic signalling and cancer metabolism. Nat Rev Cancer. 2020;20(2):74–88.

27. Fontana F, Giannitti G, Marchesi S, Limonta P. The PI3K/Akt Pathway and Glucose Metabolism: A Dangerous Liaison in Cancer. Int J Biol Sci. 2024;20(8):3113–25.

28. Dong S, Liang S, Cheng Z, Zhang X, Luo L, Li L, et al. ROS/PI3K/Akt and Wnt/â-catenin signalings activate HIF-1á-induced metabolic reprogramming to impart 5-fluorouracil resistance in colorectal cancer. J Exp Clin Cancer Res. 2022;41(1):15.

29. Nie H, Ju H, Fan J, Shi X, Cheng Y, Cang X, et al. O-GlcNAcylation of PGK1 coordinates glycolysis and TCA cycle to promote tumor growth. Nat Commun. 2020;11(1):36.

30. Jiménez-Castillo V, Illescas-Barbosa D, Zenteno E, Ávila-Curiel BX, Castañeda-Patlán MC, Robles-Flores M, et al. Increased O-GlcNAcylation promotes IGF-1 receptor/PhosphatidyI Inositol-3 kinase/Akt pathway in cervical cancer cells. Sci Rep. 2022;12(1):4464.

31. Xiang J, Chen C, Liu R, Gou D, Chang L, Deng H, et al. Gluconeogenic enzyme PCK1 deficiency promotes CHK2 O-GlcNAcylation and hepatocellular carcinoma growth upon glucose deprivation. J Clin Invest. 2021;131(8).

32. Yang Y, Xie T, Gao P, Han W, Liu Y, Wang Y. Hsa_Circ_002144 Promotes Glycolysis and Immune Escape of Breast Cancer Through miR-326/PKM Axis. Cancer Biother Radiopharm. 2024;39(10):755–69.

33. Bai X, Ali A, Lv Z, Wang N, Zhao X, Hao H, et al. Platinum complexes inhibit HER-2 enriched and triple-negative breast cancer cells metabolism to suppress growth, stemness and migration by targeting PKM/LDHA and CCND1/BCL2/ATG3 signaling pathways. Eur J Med Chem. 2021;224:113689.

34. Wang A, Zeng Y, Zhang W, Zhao J, Gao L, Li J, et al. N(6)-methyladenosine-modified SRPK1 promotes aerobic glycolysis of lung adenocarcinoma via PKM splicing. Cell Mol Biol Lett. 2024;29(1):106.

